# Immunomodulation of Pancreatic Cancer via Inhibition of SUMOylation and CD155/TIGIT Pathway

**DOI:** 10.1101/2025.02.06.636475

**Authors:** Jorge de la Torre Medina, Utsav Joshi, Himangshu Sonowal, Yixuan Kuang, Tianchen Ren, Dai-Hua Chen, Mohottige Don Neranjan Tharuka, Kim Nguyen-Ta, Hélène Gros, Zbigniew Mikulski, Yuan Chen, Rebekah R. White

## Abstract

Pancreatic ductal adenocarcinoma (PDAC) is the deadliest major cancer and has a profoundly immunosuppressive tumor microenvironment (TME). Previous studies have shown that inhibition of the E1 enzyme, which catalyzes the small ubiquitin-like modifiers (SUMO), with the small molecule TAK-981, can reprogram the TME to enhance immune activation and suppress tumor growth. We found that the CD-155/TIGIT pathway, a key regulator of immune evasion in PDAC, is influenced by SUMOylation. We hypothesized that the combination of SUMO E1 and TIGIT inhibition would synergistically induce anti-tumor immune effects. We used a clinically relevant orthotopic mouse model that consistently develops liver metastases to study this combination therapy alone and in the perioperative setting with surgical resection. The combination of SUMO E1 and TIGIT inhibition significantly prolonged survival. Complete responders exhibited protective immunity and enhanced T cell reactivity to model-specific alloantigens. Complementary immune analyses of resected tumors demonstrated that combination therapy more significantly reduces the abundance of regulatory FOXP3+CD4+ T cells than each monotherapy alone. The findings suggest that SUMO E1 inhibition enhances antibody-mediated elimination of Tregs through innate immune cells, potentially by activation of type I interferon responses. Our results highlight a mechanism to enhance the efficacy of anti-TIGIT therapy.

**Brief Summary:** SUMOylation is a post-translational modification process critical for cancer. Inhibition of SUMOylation can improve the sensitivity of pancreatic cancer to immune checkpoint inhibition.

## Introduction

Pancreatic ductal adenocarcinoma (PDAC) is the third most frequent cause of cancer-related death in the United States with an overall 5-year survival rate of only 13% (1, 2). These poor outcomes are attributable to both late presentation and treatment resistance. Cytotoxic chemotherapy can prolong survival in patients with advanced disease by a few months but is associated with significant side effects (3, 4). Even in the minority of patients with radiographically localized disease, surgical resection alone is rarely curative, since most patients have occult micrometastatic disease that later manifests as disease progression. The current standard-of-care for these patients is surgical resection combined with cytotoxic chemotherapy in the preoperative (neoadjuvant) and/or postoperative (adjuvant) settings. With these aggressive approaches, long-term survival rates following surgical resection have improved only modestly (5, 6).

Although immunotherapies targeting the PD-1 and CTLA-4 checkpoints have shown promise in other solid tumors, their results in PDAC have been disappointing (7, 8). PDAC tumors are immune-privileged and characterized by a profoundly immunosuppressive environment (9, 10) (10–12). Increased immunosuppressive cells, such as regulatory T cells (Tregs) and tumor-associated macrophages (TAMs), coupled with poor infiltration of CD8+ effector T cells into PDAC tumors and heterogeneous expression of immune checkpoints, present significant challenges for immunotherapy (13, 14). The failure of anti-PD1 and anti-CTLA-4 therapeutics in PDAC has prompted exploration of other immune checkpoints. Several recent studies have identified TIGIT (T-cell immunoreceptor with Ig and immunoreceptor tyrosine-based inhibitory motif domains) as being highly expressed in human PDAC T cells (14, 15), and associated with worse prognosis following resection (16). Pre-clinical studies have suggested that anti-TIGIT antibodies are promising as part of combinatorial approaches (17, 18).

SUMOylation is a post-translational modification involved in regulating the activity, localization, and interactions of numerous target proteins. Many proteins involved in transcription, DNA repair, and cell division are regulated by SUMOylation in cancer (19). SUMOylation enzymes are upregulated in human PDAC tumors compared to normal pancreatic tissue, and expression is inversely correlated with survival (20). Over 90% of PDAC cases are driven by mutations in K-*ras*, and amplifications of c-Myc are common and associated with metastasis (21, 22). We and others have shown that K-ras- and c-Myc-dependent oncogenesis depend on SUMOylation (23–25). SUMOylation is also involved in the regulation of immunomodulatory proteins, including Type I interferons (IFNs) and IFNγ (26, 27). This effect is similar to that of toll-like receptor agonists in that type I IFN expression activates dendritic cells for immune rejection of tumors through antitumor CD8+ T cell responses (28, 29).

Preclinical studies have highlighted the therapeutic potential of TAK-981, a SUMOylation inhibitor. The immune effects of inhibiting SUMOylation are synergistic with its direct suppressive effects on PDAC cells. TAK-981 reduced tumor burden in a subcutaneous PDAC model while increasing CD8+ T cell infiltration into tumors (20). We have developed a robust immunocompetent, orthotopic PDAC mouse model using an organoid cell line (KPC-46) that readily metastasizes to the liver (30). We have recently shown that TAK-981 improves overall survival in this model with multiple beneficial effects on the TME, including decreased immunosuppressive regulatory T cells (Tregs) (31). We also observed a significant positive correlation between expression of SUMOylation and exhaustion markers, including TIGIT, in T-cells. To explore these findings, in the current study, we hypothesized that combination of SUMOylation and TIGIT inhibition would be synergistic. Our findings demonstrate that combination therapy significantly improved overall survival in two clinically relevant metastatic PDAC models.

## Results

### SUMOylation inhibition (SUMOi) increases expression of TIGIT ligand CD155 on PDAC cells

To investigate the effects of SUMOi on the PDAC TME, we previously treated mice with orthotopic KPC-46 tumors with TAK-981 for two weeks and evaluated the transcriptional landscape within harvested tumors using single-cell RNA sequencing (31). Further analysis of this dataset revealed a significant increase in the expression of CD155 (*polio virus receptor*), the primary ligand for TIGIT, on PDAC cells following treatment with TAK-981 (Figure 1A). This upregulation of CD155 suggests enhanced susceptibility of tumor cells to immune-mediated targeting. We also observed a notable increase in CD226 expression, a co-stimulatory receptor that competes with TIGIT for binding to CD155, in the T/NK cell population (Figure 1B). These findings offer a potential mechanism by which TAK-981 treatment may modulate the TME to enhance the therapeutic efficacy of anti-TIGIT antibody (TIGITi). As further evidence that the TIGIT pathway is relevant to human PDAC, CD155 levels were significantly elevated in plasma from 3 treatment-naïve PDAC patients compared to 25 healthy subjects (Figure 1C). CD155 is similarly elevated in plasma collected from mice bearing orthotopic KPC-46 tumors compared to tumor-naïve mice (Figure 1D) with levels that correlate with tumor weight, demonstrating CD155’s potential role as a biomarker (Figure 1E).

**Figure 1.**
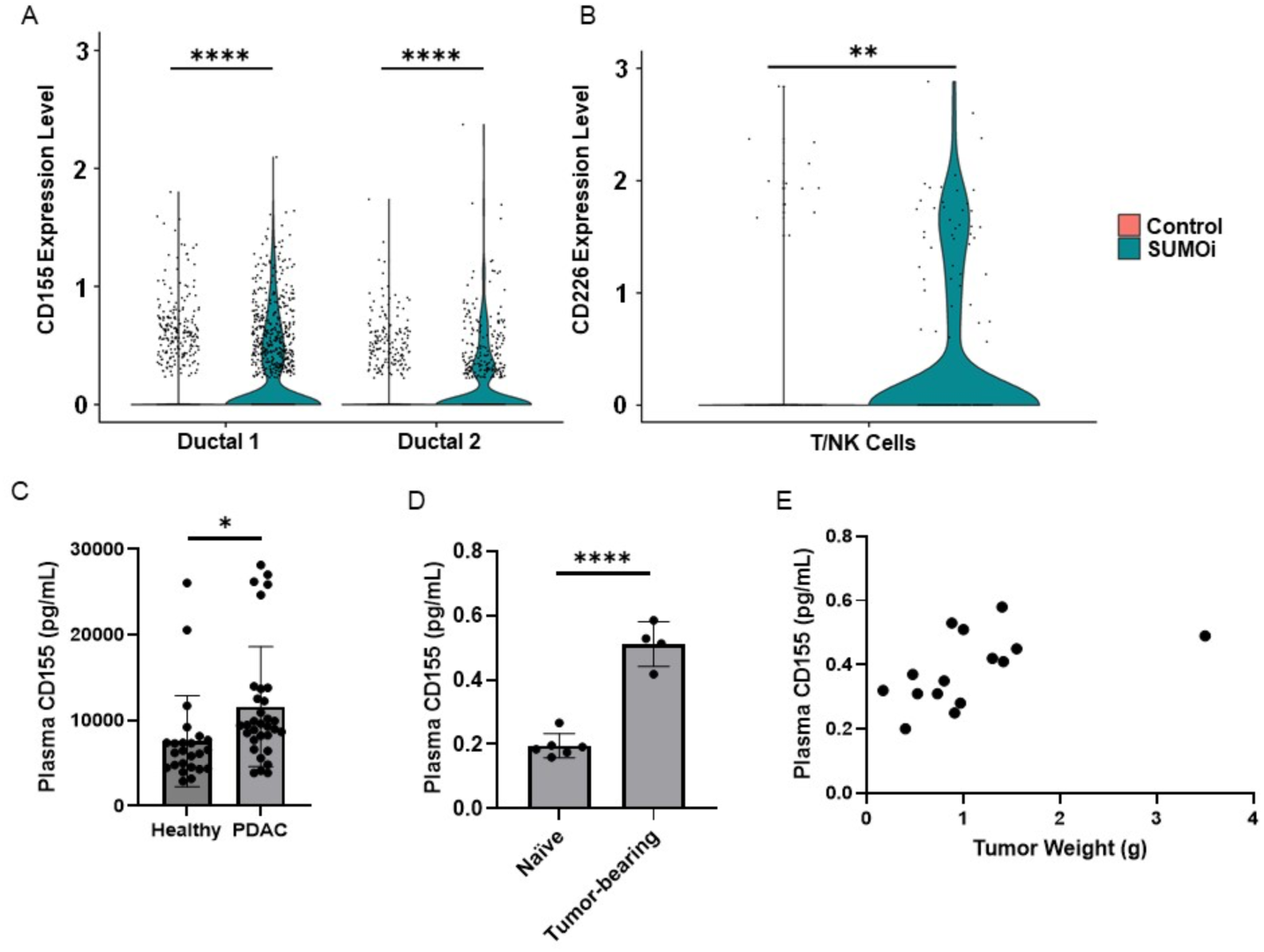
SUMOylation and TIGIT ligand CD-155 expression in PDAC. Violin plots from single cell RNA sequencing analysis of PDAC tumors showing an (A) Increase in expression of CD155 in PDAC cells and (B) increase in expression of CD226 in T and NK cells upon SUMOi treatment. (C) CD155 levels in plasma of 32 PDAC and 25 healthy volunteers (D) CD155 levels in plasma of tumor-naïve and tumor-bearing mice (E) Correlation of tumor weight with plasma CD155 levels in mice (Pearson r = 0.5, p = 0.04). Bars represent mean ± SD.

### Combined inhibition of SUMOylation and TIGIT prolongs survival in metastatic PDAC

To evaluate the potential therapeutic synergy of SUMOi and TIGITi in PDAC, we again employed the aggressively metastatic, orthotopic KPC-46 model. Once tumors reached 5–7 mm in diameter, as confirmed by ultrasound, mice were randomized into four treatment groups: SUMOi, TIGITi, SUMOi + TIGITi, or vehicle control (n=10/group). No significant differences were seen in either monotherapy group compared to the vehicle control, but there was a trend toward improved survival with the combination of SUMOi and TIGITi (p = 0.08, Figure 2A). Based on these promising results, we repeated an experiment focusing only on the combination (vs. vehicle control) and starting treatment when tumors were slightly smaller (3-5 mm). Under these conditions, the combination therapy significantly prolonged survival compared to the control group (median survival 81 vs. 42.5 days, p<0.05, Figure 2B). Furthermore, 40% of mice in the treatment group (n=4) were classified as complete responders, with no detectable disease after 2 months of observation. Most cell lines derived from KPC transgenic mice do not spontaneously metastasize to the liver when implanted orthotopically in immunocompetent mice. When mice with orthotopic tumors established from injection of two such cell lines (FC1245 and FC1199) were treated with the same combination, we did not observe significant survival benefits (Supplementary Figure), suggesting that the effects of this combination are attributable to prevention and/or control of metastatic disease.

**Figure 2.**
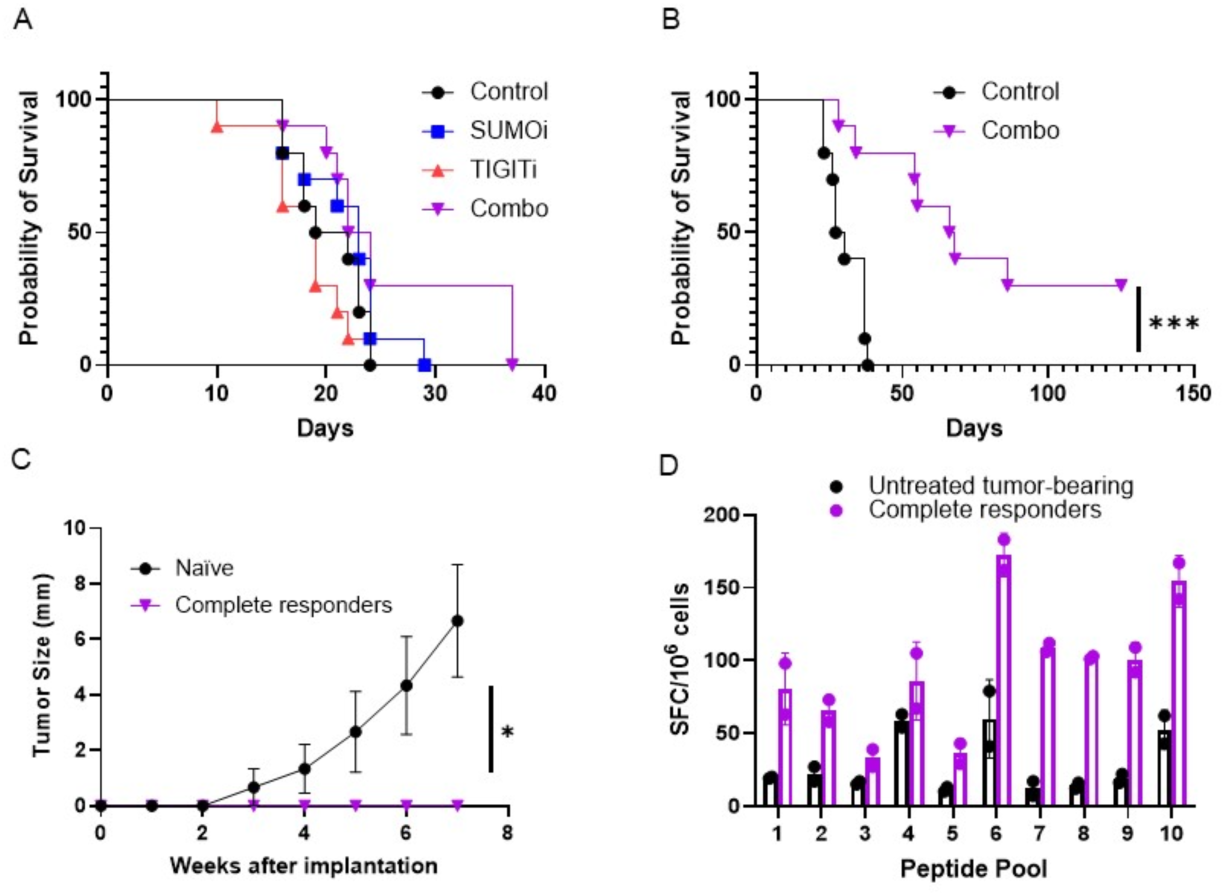
Combination of SUMOylation and anti-TIGIT improves survival and induces protective immunity. (A&B) Kaplan-Meier survival analysis showing effect of SUMOi and TIGITi, alone or in combination, in KPC46 organoid orthotopic PDAC model. (C) Complete responders rejected KPC46 organoids when implanted as subcutaneous tumors. *p<0.05, ***p=0.0004. (D) IFN-γ ELISpot assay showing greater reactivity of lymphocytes from complete responders than from untreated tumor-bearing mice to 10 pools potential peptide tumor antigens. Data represent mean ± SEM spot-forming cells (SFC)/10^6^ lymphocytes from two mice per group.

### Complete responders to combination therapy with SUMOi and TIGITi demonstrated adaptive immunity

Complete responders (n=3) and age-matched tumor-naïve controls (n=3) were rechallenged with subcutaneous injection of KPC46 tumor cells. None of the rechallenged complete responders developed tumors, while all control mice exhibited tumor growth within 2 weeks (p<0.001, Figure 2C). These findings confirm the presence of a robust anti-tumor immune response elicited by the combination therapy. We previously utilized a bioinformatic pipeline to identify and prioritize tumor-specific alloantigens (relative to the background B6/129 F1 mouse) to serve as model neoantigens for the KPC46 model (30). Peptides (two each for the top 50 candidate tumor-specific antigens) were synthesized and utilized in pools of 10 peptides in ELISpot assays to assess for neoantigen-specific T-cell responses. Following the successful rechallenge experiment, lymphocytes were collected from the spleens of complete responders and from untreated tumor-bearing mice. Complete responders exhibited significantly higher T-cell reactivity, as indicated by increased IFN-γ secretion across multiple peptide pools, compared to untreated tumor-bearing mice (Figure 2D). This heightened T-cell reactivity in complete responders suggests the formation of tumor-specific CD8+ memory T cells in response to combination therapy.

### Perioperative SUMOi and TIGITi improve survival in combination with resection

Our data suggest that the predominant effect of the combination therapy was in the immune-mediated prevention and/or control of distant metastatic disease, with relatively modest effects on primary tumor burden. We therefore evaluated this regimen in the KPC46 model in combination with resection of the primary tumor to model neoadjuvant and adjuvant therapy given with curative intent to patients with radiographically-localized disease. Neoadjuvant treatment commenced 7 days post-implantation, followed by resection then matched adjuvant therapy (Figure 3A). Combination therapy, but neither monotherapy, significantly improved overall survival (Figure 3B). One mouse in the combination group and one in the perioperative SUMOi group were considered “cured” with no evidence of disease recurrence at over 100 days All of the remaining mice died with gross evidence of metastatic disease. There was only a trend toward decreased primary tumor weight in the combination group after two weeks of therapy (Figure 3C), supporting our theory that the survival benefit seen with this combination is attributable to control of metastatic disease. We also observed a significant inverse relationship between resected tumor weight and survival (r = −0.58, p<0.0001). This illustrates the validity of this model for studying human PDAC, as tumors consistently metastasize to the liver at a relatively small size, before mice die from their local tumor burden.

**Figure 3.**
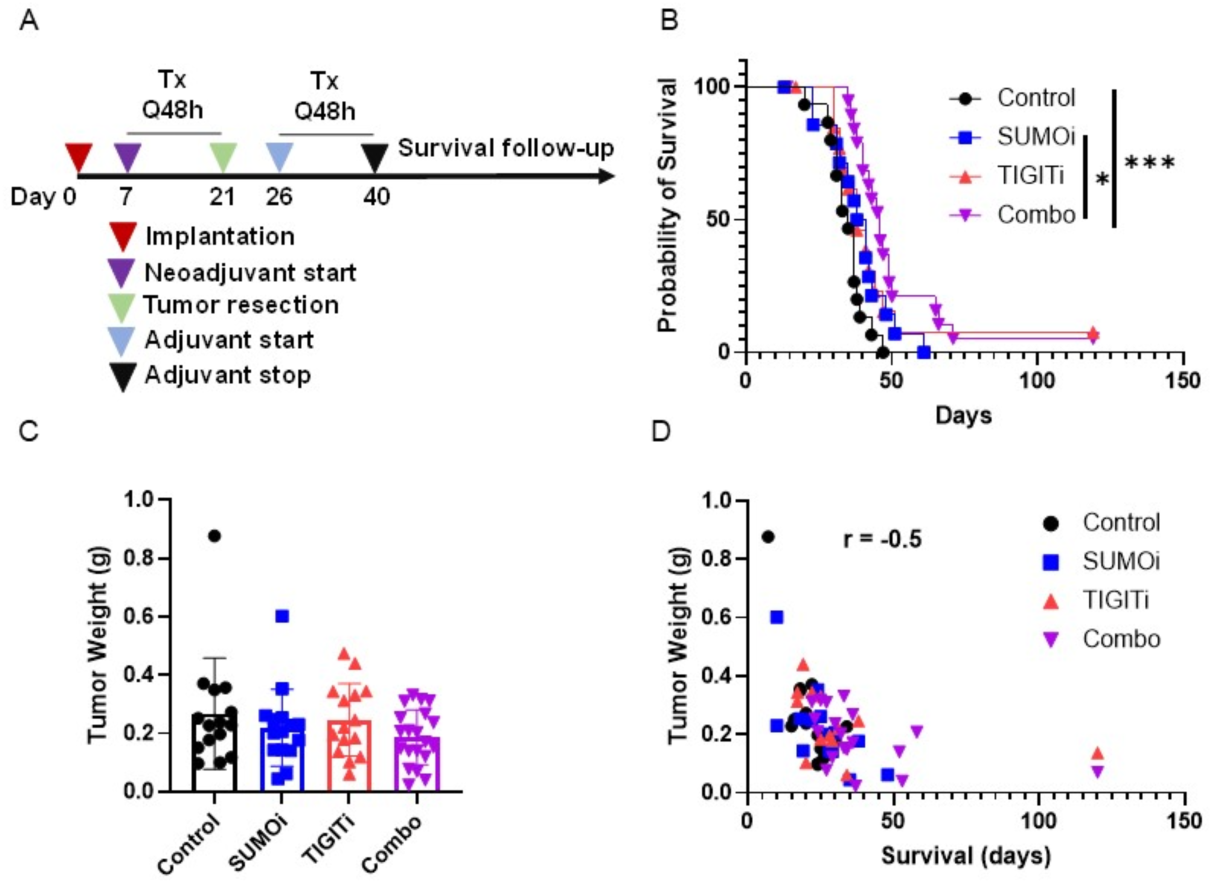
Combination neoadjuvant therapy followed by resection and adjuvant therapy with SUMOylation and anti-TIGIT improves survival in orthotopic KPC46 orthotopic pancreatic organoid model. (A) Study design (B) Kaplan-Meier survival analysis of tumor-bearing mice treated with SUMOi and TIGITi, alone or in combination. Statistical significance was determined by the log-rank test. *p<0.001 (C) Weight of KPC46 orthotopic organoid tumors harvested 2 weeks after neoadjuvant therapy. (D) Correlation of tumor size on day of resection and survival time in mice (Pearson r = −0.05, p<0.0001).

### Combined Inhibition of SUMOylation and TIGIT decreases tumor-infiltrating T regulatory cells

Flow cytometric analysis was performed for tumor infiltrating T cells and myeloid cells. No significant changes were observed for the myeloid or CD8^+^ T cell populations. However, we found that CD4^+^ T cells were significantly decreased in both the SUMOi monotherapy group and the SUMOi + TIGITi combination group (Figure 4A). Further analysis shows that the decrease in CD4^+^ T cells is mainly due to the decrease of Treg populations (CD4^+^FOXP3^+^), which were significantly reduced in the combination group (Figure 4B). Tumor-infiltrated Th1 cells were not significantly reduced in any treatment groups (Figure 4C). In addition to the reduction in Treg populations, co-inhibitory receptors on Tregs, such as PD-1, TIM-3, and TIGIT, were significantly decreased in the combination treatment group (Figure 4E, 4F, 4G). Taken together, the data suggest that, within the tumor microenvironment, SUMOi and TIGITi synergize in reducing Treg populations and the suppressive receptors of Tregs required for their immune suppressive functions.

**Figure 4.**
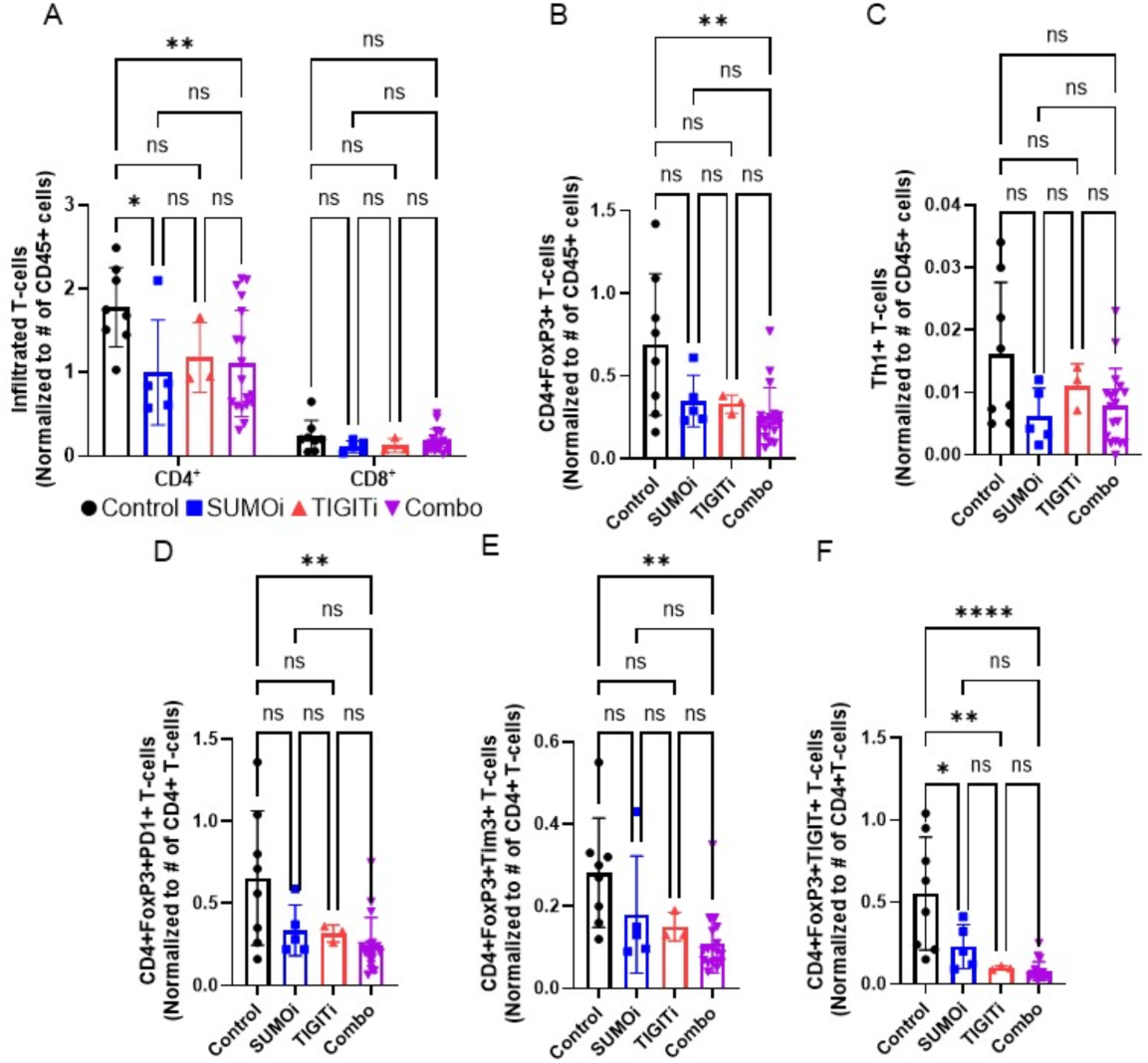
Combined Inhibition of SUMOylation and TIGIT decreases tumor-infiltrating T regulatory cells and inhibits expression of T-cell exhaustion makers. Flow cytometry analysis of tumors analyzing the number of (A) Tumor-infiltrating CD4+ and CD8+ T-cells (B) CD4+FoxP3+ T-cells (C) Th1+ T-bet cells (D) CD4+FoxP3+PD1+ T-cells (E) CD4+FoxP3+Tim3+ T-cells (F) CD4+FoxP3+TIGIT+ T-cells. Data are mean ± SD. **<0.001, ***p<0.0001, ****p<0.00001.

## Discussion

The vast majority of patients diagnosed with PDAC will either present with or develop distant metastatic disease. Cytotoxic chemotherapy is rarely curative in PDAC, and immune checkpoint inhibitors have shown limited efficacy in PDAC due to factors such as poor infiltration of CD8+ effector T cells and the dominance of immunosuppressive cells, including T regs, in the TME. Innovative strategies to overcome the PDAC TME are necessary to meaningfully improve outcomes in this disease.

SUMOylation is a ubiquitous post-translational modification process critical for modulating protein activity by controlling their interacting partners and localization through a SUMO-interacting motif (32). PDAC tumor samples exhibit upregulated SUMOylation enzymes, which correlate with poor patient outcomes (19). Inhibition of SUMOylation has been shown to inhibit tumor cell proliferation *in vitro* and in xenograft models in immunocompromised mice (20, 33, 34), confirming its direct effects on cancers cells. However, we have shown that the effects of SUMO inhibition appear to be mainly attributable to its many effects on immune cells in an orthotopic PDAC model (31). SUMOylation is known to be required for T reg proliferation and function through stabilization of transcription factor interferon regulatory factor 4, IRF4 (35). In our prior study, SUMO inhibition significantly decreased Treg abundance in the PDAC TME and increased activation of cytotoxic T cells, macrophages, and NK cells (36). Expression of SUMOylation enzymes was also positively correlated with an “exhaustion score” reflecting multiple exhaustion markers, including TIGIT (36).

Our focus on TIGIT in the current study was driven by multiple studies by others showing that TIGIT and CD155 are more highly expressed in PDAC than PD1 and PDL1 (13–15). This was corroborated by our finding that CD155 levels were significantly increased in PDAC patient plasma and in mice bearing KPC-46 tumors. Increased CD155 expression in human PDAC tissue has previously been associated with worse outcomes following resection (37), and we found that CD155 expression was positively correlated with tumor size in our mouse model. SUMO inhibition led to an increase in CD155 expression on PDAC cells and increased expression of TIGIT’s competitor co-stimulatory receptor CD226 on T and NK cells in our model. High expression of CD226 in CD8+ T cells has been associated with improved response to anti-TIGIT therapy (38), and together these findings led us to hypothesize that SUMO inhibition would enhance the efficacy of TIGIT inhibition We found that the combination of SUMOi and TIGITi prolonged survival in an aggressively metastatic orthotopic PDAC model but with only modest effects on primary tumor growth. We utilized a model of perioperative therapy to demonstrate the relevance of this combination therapy to the many PDAC patients with radiographically localized disease but occult distant micrometastases. A few studies have utilized survival distal pancreatectomy in xenograft mouse models to study recurrence (39, 40). However, most immunocompetent mouse PDAC models— aside from their parent genetically-engineered mouse models—do not develop spontaneous liver metastases. This organoid line, derived from a liver metastasis in a KPC mouse, facilitates the study of arguably this most important feature of human PDAC.

We found that SUMOi synergizes with TIGITi to reduce the abundance and inhibitory activity of Tregs. This is possibly due to enhanced antibody-dependent cellular cytotoxicity (ADCC) or antibody-dependent cellular phagocytosis (ADCP) by SUMOi to eliminate Tregs (Figure 5). Reduction of Tregs by SUMOi is synergistic with direct inhibition of Tregs by SUMOi. In addition, anti-TIGIT blocks the inhibitory signal in cytotoxic T lymphocytes, thereby enhancing the interaction between the co-stimulatory receptor CD226 and CD155 on tumor cells, both of which are both increased by SUMOi (Figure 1A, 1B, and 5). Although we did not see a significant increase in the number of infiltrating CD8+ effector cells, which are sparse in this model, we did see evidence of T-cell activity. Our rechallenge experiments revealed the development of protective immunity in complete responders. Increased T-cell reactivity to previously identified model-specific tumor antigens (30) observed in our ELISpot assay supports that this immunity is T-cell-mediated. These observations align with previous studies showing that SUMO inhibition can enhance antigen presentation and CD8+ T cell infiltration in lymphoma and colon cancer mouse models (41).

**Figure 5.**
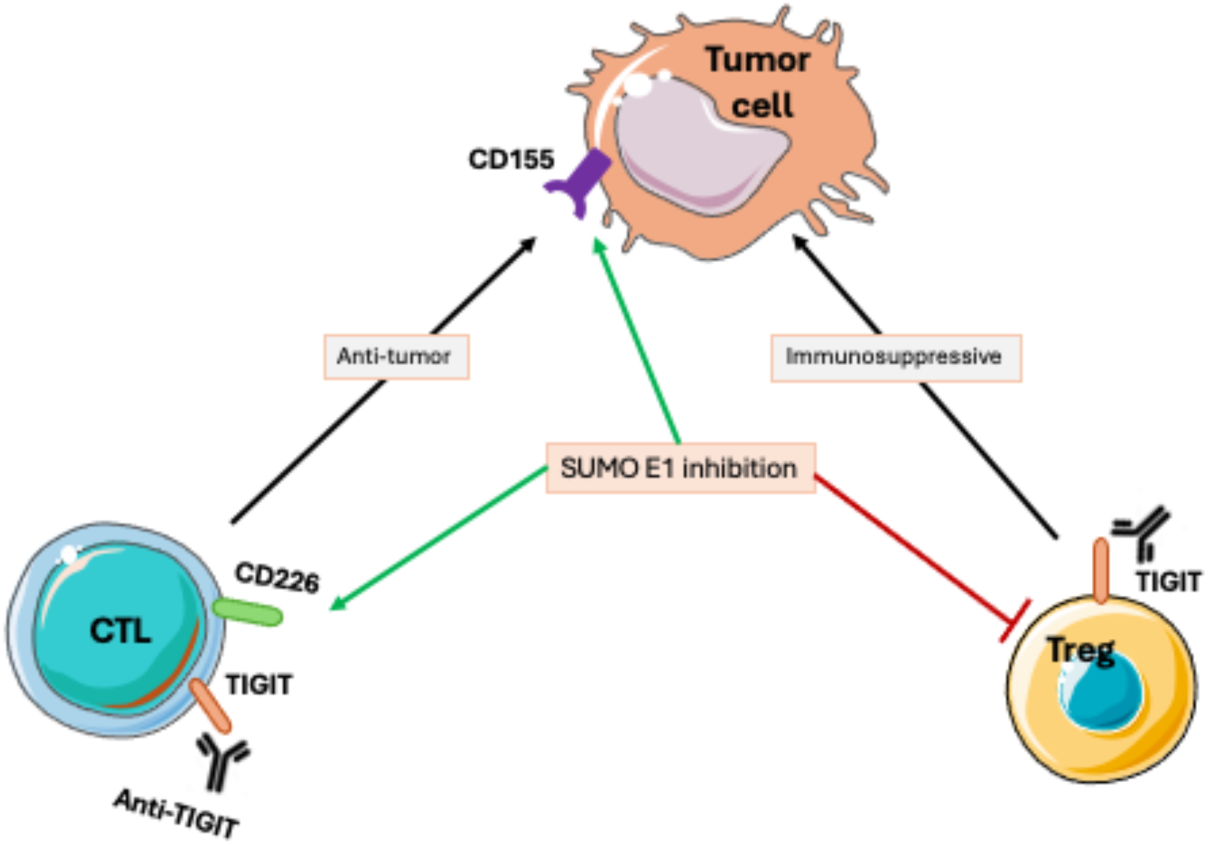
Proposed mechanism of synergy. SUMO E1 inhibition has direct inhibitory effects on immunosuppressive T regulatory (Treg) cells. In parallel, SUMO E1 inhibition increases expression of TIGIT ligand CD155 on tumor cells and CD226 on cytotoxic T lymphocytes (CTLs), promoting anti-tumor immunity.

Despite these promising results, several limitations must be acknowledged. The KPC-46 model is highly aggressive and variable in its growth rate. We observed significant differences in complete response rates over repeated experiments, ranging from 0 to 40%, which we believe was attributable to the timing of treatment initiation. The relatively low cure rate in our perioperative therapy experiment is a testament to the aggressiveness of this model. Furthermore, our model, while robust, does not reflect the heterogeneity of human PDAC. Mouse models driven by K-ras and P53 mutations have been criticized for being poorly immunogenic (42). However, the mixed genetic background of tumors derived from KPC mice results in the generation of single nucleotide polymorphisms relative to the host mouse that can serve as model antigens capable of generating quantifiable T-cell reactivity (30, 43).

The frequent co-expression of TIGIT with PD1 on immune cells has provided rationale for clinical studies of combined approaches in other tumor types. Although several early phase studies showed encouraging activity, two recent randomized studies of standard therapy including anti-PDL1 antibody with or without anti-TIGIT antibody in non-small cell lung cancer were negative (44, 45). A follow-up biomarker study, however, demonstrated that tumors with high baseline Tregs and macrophages did demonstrate improved responses with the addition of anti-TIGIT therapy (46), which bodes well for PDAC.

In conclusion, our study demonstrates the potential for SUMO inhibition to modulate the PDAC TME and enhance the efficacy of TIGIT inhibition. There are multiple TIGIT inhibitors that are currently in preclinical development or already in clinical trials (47). Another approach to inhibit the SUMO E1 enzyme has been identified through a covalent, allosteric mechanism that likely offers prolonged pharmacodynamic effects (48, 49). An orally bioavailable SUMO E1 inhibitor has been developed based on this mechanism (50). TAK-981 has had a favorable toxicity and efficacy profile in several early phase trials, paving the way to clinical development for other inhibitors of SUMOylation enzymes that may offer improved efficacy. We believe that the combination of SUMO E1 and TIGIT inhibition is ready for translation with parallel exploration of potential predictive biomarkers.

## MATERIALS AND METHODS

### Sex as a Biological Variable

This study included both male and female PDAC cell lines in sex-matched mice as well as human plasma samples from men and women.

### Single-cell RNA sequencing (scRNAseq)

The single cell RNA sequencing data were processed via the methods described previously (31). The gene expression data was obtained from the Seurat analysis and plotted using R packages Seurat and ggplot2. The statistical analysis was done using the stat_compare_means function from the R package ggpubr.

### Plasma Analysis for CD155

Sandwich ELISA was used to quantify human CD226/DNAM-1(Biotechne R&D Systems, DY666-05) in human plasma and mouse CD155/PVR (Biotechne R&D Systems) in mouse plasma. De-identified plasma samples from treatment-naive patients with PDAC were obtained through the UC San Diego Moores Cancer Center Biorepository (IRB protocol #181755). Blood was collected into lithium heparin tubes (green-top Becton BD Vacutainer). Samples from healthy volunteer subjects were also collected under IRB protocol #100556. Additional normal human plasma samples were obtained from Innovative Research. Mouse plasma samples were collected at the time of euthanasia from untreated mice bearing orthotopic KPC46 tumors and tumor-naive mice by cardiac puncture. The samples were stored at −80 °C, thawed at 4°C, and handled on ice for the assay. ELISA assays were performed following the manufacturers’ protocols. Optical density was determined using a SpectraMax iD3 (Molecular Devices) plate reader at 450nm. Blank readings were subtracted from each well. The data were analyzed using a non-linear, sigmoidal model with least squares regression, and unknown concentration values were interpolated from a standard curve with 95% confidence in GraphPad Prism software.

### Cell Lines and Organoids

KPC46 organoids were derived from a male *Kras^LSL-G12D^p53^LSL-R172H^-Pdx1-Cre* (KPC) mouse with a B6/129 F1 hybrid background (gift of Andrew Lowy, UC San Diego (51)) and were maintained in RPMI supplemented with 5% FBS, 1x penicillin-streptomycin, 1 mM Glutamax, 1 mM sodium pyruvate, 1x nonessential amino acids, 1x Fungizone, 5 µg/mL insulin, 1.4 µM hydrocortisone, 10 ng/mL EGF, and 10.5 µM ROCK inhibitor. FC1199 (male) and FC1245 (female) murine pancreatic cancer cells were isolated from KPC transgenic mice on a homogenous C57BL6 background and were shared with Dr. Andrew Lowy by Dr. David Tuveson at Cold Springs Harbor Laboratory (52). Cells were maintained in DMEM/Hams F12 50:50 (Corning, 10-090-CV) supplemented with 10% FBS and 1x penicillin streptomycin.

### Animals

All animal experiments were approved by the Institutional Animal Care and Use Committee (IACUC) and conducted in an AAALAC-accredited facility at UC San Diego (UCSD). 6–8-week-old wild-type C57BL/6 (male/female) and B6/129 F1 hybrid mice (male) were purchased from Jackson Laboratories (Bar Harbor, ME). All mice were housed in pathogen-free conditions with ad libitum access to food and water. To establish the orthotopic organoid model, organoids were enzymatically dissociated from Matrigel™ (Corning) domes and sheared into single organoid suspensions using a 23G needle. Organoids were treated with Cell Recovery Solution (Corning) and TrypLE Express (Thermo Fisher Scientific) for 1 hour at 37°C to generate a single-cell suspension. Cells were counted, centrifuged (2000 RPM, 15 min, 4°C), and resuspended in growth factor-depleted Matrigel at a density of 2.5 × 10⁶ cells/mL. A 20 µL aliquot of this suspension was orthotopically injected into the pancreatic tail of 6-to-8 week-old male B6/129 F1 mice via laparotomy. FC1199 and FC1245 were trypsinized, counted and resuspended at a density of 0.25 x 10^6^ cells/mL in Matrigel and 20 µL was injected into the pancreatic tail of male and female C57BL/6, respectively, via laparotomy. Mice received 1 mg/kg buprenorphine for analgesia prior to surgery.

### Therapeutic Experiments in Mice

Tumor engraftment after orthotopic implantation was monitored using handheld ultrasound (SonoQue L5P). Once tumors reached 3–7 mm in diameter, mice were randomized into treatment groups as described for respective experiments in Results. TAK-981 used in the study was provided by Takeda Development Center Americas, Inc., or purchased from Chemietek (Catalog number: CT-TAK981). TAK-981 purchased from Chemietek was verified by NMR and LC-MS and by enzymatic inhibition assay. TAK-981 was delivered at a dose of 15 mg/kg, dissolved in ultrapure DMSO and resuspended in 5% Dextrose and 20% Kolliphor (Sigma Aldrich, 30906). Functional grade anti-mouse TIGIT antibody (Leinco Technologies, T735, Clone 1G9) was diluted in diluent buffer (Leinco Technologies, D230) and delivered at a dose of 100 µg/mice. In most of the survival studies, mice were treated for 3 weeks and then monitored until death or euthanasia due to meeting humane endpoints. In the surgical resection experiment, neoadjuvant treatment was started 7 days after orthotopic tumor implantation. After 2 weeks of treatment, mice underwent a second laparotomy. A brief abdominal survey was performed to confirm the absence of gross metastatic disease. Distal pancreatectomy was then performed to include the tumor with a grossly negative margin, using a 5 mm titanium clip (angioOCCLUDE, Medline Industries) at the base of the normal pancreas after dividing the gland with electrocautery. Five days post-resection, mice resumed treatment with the same regimen given preoperatively for an additional 2-week course of adjuvant therapy to complete a total of 4 weeks of treatment.

### Rechallenge Experiment

KPC46 organoids (20 µL at a density of 2.5 × 10⁶ cells/mL), suspended in Matrigel, were injected into right flank of complete responders or age-matched male B6/129 F1J mice. Mice were monitored daily for tumor presence. Tumor sizes were measured using Vernier calipers.

### ELISpot assay

Peptides (20 amino acids in length) harboring potential antigens previously identified were synthesized by TC peptide Labs (San Diego, CA) and separated into peptide pools of 10 peptides each as described previously (30). Lymphocytes were isolated from the spleens of complete responders to treatment and untreated tumor-bearing control mice, and bone marrow-derived dendritic cells (BMDCs) were isolated from donor B6/129F1J mice as described previously (30). ELISPOT kit (interleukin-2 and IFN-γ) was purchased from Immunospot (Cleveland, OH), and the assay was performed according to manufacturer instruction. Briefly pre-wet high binding PVDF filter plates were coated with capture IFN-y antibody. BMDCs were incubated with 5 µg/mL of mutant peptide pools to facilitate antigen presentation for 24 h at 37°C. 2×10^5^ lymphocytes isolated from control and rechallenged mice were incubated with 5 µg/mL of the different mutant peptide pools to which 20,000 activated BMDCs were added. A 5 µg/mL of concanavalin A (Sigma-Aldrich) was used as the positive stimulus control, and no peptide wells were used as the negative control. IFNγ secretion was assessed with biotinylated anti-mouse IFNγ and imaged using an ELISPOT reader (AID Diagnostika).

### Flow Cytometry

Tumors were harvested and dissociated into single cell suspension using a tumor dissociation kit (Milteni Biotec, 130-096-730) following manufacturer’s protocol. After obtaining single cell suspension, RBC lysis was carried out, counted and washed with cell staining buffer (2% FBS, 5 mM EDTA, and Dulbecco’s Phosphate-Buffered Saline). The cell suspensions were first stained with LIVE/DEAD Fixable Blue Dead Cell Stain Kit (ThermoFisher, L23105) at 4 °C for 30 minutes, protected from light. The cells were then washed 3x with cell staining buffer and processed for cell surface staining for 30 min at 4 °C protected from light with the following anti-mouse antibodies: CD45 (Cytek, 30-F11), CD3 (BD biosciences, 563565), Viability blue Live/Dead (ThermoFisher, L23105), CD8 (ThermoFisher, 368-0081-82), CD4 (Bio-Rad, MCA2691SBUV605), CD69 (Bio-Legend, 104530), CD62L (Bio-Legend, 104450), CD44 (ThermoFisher, 365-044-82), CD127 (Bio-Legend, 158214), CD25 (Bio-Legend, 102048), TIM3 (Bio-Legend, 134010), PD1 (BD biosciences, 568603), CD11c (Bio-Legend, 117368), CD103 (BD biosciences, 566118), TIGIT (BD biosciences, 565270). After cell surface staining, the cells were first washed 3x with cell staining buffer and then fixed at 4% formaldehyde at 4 °C for 30 minutes, protected from light. A wash process was performed after fixation. For intracellular staining, the post-surface-staining cells were permeabilized and fixed with Foxp3/Transcription Factor Staining Buffer Set (ThermoFisher, 00-5523-00) following manufacturer’s recommended procedures. Anti-mouse intracellular antibodies were TCF1/7 (Cell signaling, 14456S), FOXP3 (ThermoFisher, MA5-18160), Tbet (BD biosciences, 568167), TOX (BD biosciences, 570193). Post-intracellular staining cells were washed 2x with permeabilization buffer. Samples were acquired using Cytek Aurora 5L full spectrum flow cytometer and analyzed using FlowJo TM v10 Software.

### Statistical Analysis

Results are expressed as means ± standard error of the mean (SEM) or mean ± standard deviation (SD). Statistical difference between groups was calculated either using the student’s *t* test or ANOVA with post-hoc for multiple comparisons depending on the data, using GraphPad Prism software. Pearson correlation analysis was used to evaluate the relationship between two parameters. Survival data between groups was analyzed using the Kaplan-Meier method. Statistical significance was defined as a p-value of ≤ 0.05.

### Data Availability Statement

All data relevant to the study are included in the article and associated supplementary files. All other raw data are available upon reasonable request from the corresponding author.

## Author Contributions

The authors’ contributions were as follows: designing research studies (JDLT, UJ, ZM, YC, RRW), conducting experiments (JDLT, UJ, HS, KN, HG), acquiring data (YK, LR, DC, MDNT), analyzing data (JDLT, UJ, HS, YK, YC, RRW), providing reagents (YC), and writing the manuscript (JDLT, HS, YK, KN, YC, RRW).

## Acknowledgments

Cell lines and valuable guidance on mouse models were provided by UCSD colleagues Andrew Lowy and Herve Tiriac. This publication includes data generated at the UC San Diego IGM Genomics Center utilizing an Illumina NovaSeq 6000 that was purchased with funding from a National Institutes of Health SIG grant (#S10 OD026929). We also thank the Biorepository and Tissue Technology Core facility at Moores Cancer Center for assistance in collection of PDAC patient plasma samples. The study described here was supported by the following grants: NIH R01 CA254268 (RRW) with a Research Supplement to Promote Diversity in Health-Related Research (JDLT), NIH R01CA265410 (YC), R01CA212119 (YC), and Pancreatic Cancer Action Network Translational Research Grant (19-65, YC).

## References

1. Surveillance, Epidemiology, and End Results (SEER) Program (www.seer.cancer.gov): Cancer Stat Facts [Internet]. [cited January 3, 2025]. Available from: https://seer.cancer.gov/statfacts/html/pancreas.html.

2. Siegel RL, Kratzer TB, Giaquinto AN, Sung H, Jemal A. Cancer statistics, 2025. CA Cancer J Clin. 2025.

3. Conroy T, Desseigne F, Ychou M, Bouche O, Guimbaud R, Becouarn Y, et al. FOLFIRINOX versus gemcitabine for metastatic pancreatic cancer. The New England journal of medicine. 2011;364(19):1817–25.

4. Von Hoff DD, Ervin T, Arena FP, Chiorean EG, Infante J, Moore M, et al. Increased survival in pancreatic cancer with nab-paclitaxel plus gemcitabine. The New England journal of medicine. 2013;369(18):1691–703.

5. Conroy T, Hammel P, Hebbar M, Ben Abdelghani M, Wei AC, Raoul JL, et al. FOLFIRINOX or Gemcitabine as Adjuvant Therapy for Pancreatic Cancer. The New England journal of medicine. 2018;379(25):2395–406.

6. Ahmad SA, Duong M, Sohal DPS, Gandhi NS, Beg MS, Wang-Gillam A, et al. Surgical Outcome Results From SWOG S1505: A Randomized Clinical Trial of mFOLFIRINOX Versus Gemcitabine/Nab-paclitaxel for Perioperative Treatment of Resectable Pancreatic Ductal Adenocarcinoma. Annals of surgery. 2020;272(3):481–6.

7. Royal RE, Levy C, Turner K, Mathur A, Hughes M, Kammula US, et al. Phase 2 trial of single agent Ipilimumab (anti-CTLA-4) for locally advanced or metastatic pancreatic adenocarcinoma. Journal of immunotherapy. 2010;33(8):828–33.

8. O’Reilly EM, Oh DY, Dhani N, Renouf DJ, Lee MA, Sun W, et al. Durvalumab With or Without Tremelimumab for Patients With Metastatic Pancreatic Ductal Adenocarcinoma: A Phase 2 Randomized Clinical Trial. JAMA Oncol. 2019;5(10):1431–8.

9. Balachandran VP, Luksza M, Zhao JN, Makarov V, Moral JA, Remark R, et al. Identification of unique neoantigen qualities in long-term survivors of pancreatic cancer. Nature. 2017;551(7681):512-6.

10. Beatty GL, Werba G, Lyssiotis CA, Simeone DM. The biological underpinnings of therapeutic resistance in pancreatic cancer. Genes Dev. 2021;35(13-14):940–62.

11. Clark CE, Hingorani SR, Mick R, Combs C, Tuveson DA, Vonderheide RH. Dynamics of the immune reaction to pancreatic cancer from inception to invasion. Cancer Res. 2007;67(19):9518–27.

12. Binnewies M, Roberts EW, Kersten K, Chan V, Fearon DF, Merad M, et al. Understanding the tumor immune microenvironment (TIME) for effective therapy. Nat Med. 2018;24(5):541–50.

13. Liudahl SM, Betts CB, Sivagnanam S, Morales-Oyarvide V, da Silva A, Yuan C, et al. Leukocyte Heterogeneity in Pancreatic Ductal Adenocarcinoma: Phenotypic and Spatial Features Associated with Clinical Outcome. Cancer Discov. 2021;11(8):2014–31.

14. Werba G, Weissinger D, Kawaler EA, Zhao E, Kalfakakou D, Dhara S, et al. Single-cell RNA sequencing reveals the effects of chemotherapy on human pancreatic adenocarcinoma and its tumor microenvironment. Nat Commun. 2023;14(1):797.

15. Steele NG, Carpenter ES, Kemp SB, Sirihorachai VR, The S, Delrosario L, et al. Multimodal Mapping of the Tumor and Peripheral Blood Immune Landscape in Human Pancreatic Cancer. Nat Cancer. 2020;1(11):1097–112.

16. Heiduk M, Klimova A, Reiche C, Digomann D, Beer C, Aust DE, et al. TIGIT Expression Delineates T-cell Populations with Distinct Functional and Prognostic Impact in Pancreatic Cancer. Clin Cancer Res. 2023;29(14):2638–50.

17. Freed-Pastor WA, Lambert LJ, Ely ZA, Pattada NB, Bhutkar A, Eng G, et al. The CD155/TIGIT axis promotes and maintains immune evasion in neoantigen-expressing pancreatic cancer. Cancer cell. 2021;39(10):1342–60 e14.

18. Peng H, Li L, Zuo C, Chen MY, Zhang X, Myers NB, et al. Combination TIGIT/PD-1 blockade enhances the efficacy of neoantigen vaccines in a model of pancreatic cancer. Front Immunol. 2022;13:1039226.

19. Schneeweis C, Hassan Z, Schick M, Keller U, Schneider G. The SUMO pathway in pancreatic cancer: insights and inhibition. Br J Cancer. 2021;124(3):531–8.

20. Kumar S, Schoonderwoerd MJA, Kroonen JS, de Graaf IJ, Sluijter M, Ruano D, et al. Targeting pancreatic cancer by TAK-981: a SUMOylation inhibitor that activates the immune system and blocks cancer cell cycle progression in a preclinical model. Gut. 2022.

21. Witkiewicz AK, McMillan EA, Balaji U, Baek G, Lin WC, Mansour J, et al. Whole-exome sequencing of pancreatic cancer defines genetic diversity and therapeutic targets. Nat Commun. 2015;6:6744.

22. Maddipati R, Norgard RJ, Baslan T, Rathi KS, Zhang A, Saeid A, et al. MYC Levels Regulate Metastatic Heterogeneity in Pancreatic Adenocarcinoma. Cancer Discov. 2022;12(2):542–61.

23. Kessler JD, Kahle KT, Sun T, Meerbrey KL, Schlabach MR, Schmitt EM, et al. A SUMOylation-dependent transcriptional subprogram is required for Myc-driven tumorigenesis. Science. 2012;335(6066):348-53.

24. Li YJ, Du L, Aldana-Masangkay G, Wang X, Urak R, Forman SJ, et al. Regulation of miR-34b/c-targeted gene expression program by SUMOylation. Nucleic Acids Res. 2018;46(14):7108–23.

25. Yu B, Swatkoski S, Holly A, Lee LC, Giroux V, Lee CS, et al. Oncogenesis driven by the Ras/Raf pathway requires the SUMO E2 ligase Ubc9. Proceedings of the National Academy of Sciences of the United States of America. 2015;112(14):E1724–33.

26. Decque A, Joffre O, Magalhaes JG, Cossec JC, Blecher-Gonen R, Lapaquette P, et al. Sumoylation coordinates the repression of inflammatory and anti-viral gene-expression programs during innate sensing. Nat Immunol. 2016;17(2):140–9.

27. Crowl JT, Stetson DB. SUMO2 and SUMO3 redundantly prevent a noncanonical type I interferon response. Proceedings of the National Academy of Sciences of the United States of America. 2018;115(26):6798–803.

28. Fuertes MB, Kacha AK, Kline J, Woo SR, Kranz DM, Murphy KM, et al. Host type I IFN signals are required for antitumor CD8+ T cell responses through CD8{alpha}+ dendritic cells. J Exp Med. 2011;208(10):2005–16.

29. Diamond MS, Kinder M, Matsushita H, Mashayekhi M, Dunn GP, Archambault JM, et al. Type I interferon is selectively required by dendritic cells for immune rejection of tumors. J Exp Med. 2011;208(10):1989–2003.

30. Shankara Narayanan JS, Hayashi T, Erdem S, McArdle S, Tiriac H, Ray P, et al. Treatment of pancreatic cancer with irreversible electroporation and intratumoral CD40 antibody stimulates systemic immune responses that inhibit liver metastasis in an orthotopic model. J Immunother Cancer. 2023;11(1).

31. Erdem S, Lee HJ, Shankara Narayanan JSN, Tharuka MDN, De la Torre J, Ren T, et al. Inhibition of SUMOylation Induces Adaptive Antitumor Immunity against Pancreatic Cancer through Multiple Effects on the Tumor Microenvironment. Mol Cancer Ther. 2024;23(11):1597–612.

32. Song J, Durrin LK, Wilkinson TA, Krontiris TG, Chen Y. Identification of a SUMO-binding motif that recognizes SUMO-modified proteins. Proceedings of the National Academy of Sciences of the United States of America. 2004;101(40):14373–8.

33. Langston SP, Grossman S, England D, Afroze R, Bence N, Bowman D, et al. Discovery of TAK-981, a First-in-Class Inhibitor of SUMO-Activating Enzyme for the Treatment of Cancer. Journal of medicinal chemistry. 2021;64(5):2501–20.

34. Kotani H, Oshima H, Boucher JC, Yamano T, Sakaguchi H, Sato S, et al. Dual inhibition of SUMOylation and MEK conquers MYC-expressing KRAS-mutant cancers by accumulating DNA damage. J Biomed Sci. 2024;31(1):68.

35. Ding X, Wang A, Ma X, Demarque M, Jin W, Xin H, et al. Protein SUMOylation Is Required for Regulatory T Cell Expansion and Function. Cell Rep. 2016;16(4):1055–66.

36. Erdem S, Lee H, Narayanan J, N. T, De la Torre J, Ren T, et al. Inhibition of SUMOylation Induces Adaptive Anti-Tumor Immunity Against Pancreatic Cancer through Multiple Effects on the Tumor Microenvironment. Molecular cancer therapeutics. 2024;in press.

37. Zhang D, Liu J, Zheng M, Meng C, Liao J. Prognostic and clinicopathological significance of CD155 expression in cancer patients: a meta-analysis. World J Surg Oncol. 2022;20(1):351.

38. Jin HS, Ko M, Choi DS, Kim JH, Lee DH, Kang SH, et al. CD226(hi)CD8(+) T Cells Are a Prerequisite for Anti-TIGIT Immunotherapy. Cancer Immunol Res. 2020;8(7):912–25.

39. Pang TCY, Xu Z, Mekapogu AR, Pothula S, Becker T, Corley S, et al. HGF/c-Met Inhibition as Adjuvant Therapy Improves Outcomes in an Orthotopic Mouse Model of Pancreatic Cancer. Cancers (Basel). 2021;13(11).

40. Ni X, Yang J, Li M. Imaging-guided curative surgical resection of pancreatic cancer in a xenograft mouse model. Cancer Lett. 2012;324(2):179–85.

41. Lightcap ES, Yu P, Grossman S, Song K, Khattar M, Xega K, et al. A small-molecule SUMOylation inhibitor activates antitumor immune responses and potentiates immune therapies in preclinical models. Sci Transl Med. 2021;13(611):eaba7791.

42. Evans RA, Diamond MS, Rech AJ, Chao T, Richardson MW, Lin JH, et al. Lack of immunoediting in murine pancreatic cancer reversed with neoantigen. JCI Insight. 2016;1(14).

43. Narayanan JSS, Ray P, Hayashi T, Whisenant TC, Vicente D, Carson DA, et al. Irreversible Electroporation Combined with Checkpoint Blockade and TLR7 Stimulation Induces Antitumor Immunity in a Murine Pancreatic Cancer Model. Cancer Immunol Res. 2019;7(10):1714–26.

44. Cho BC, Abreu DR, Hussein M, Cobo M, Patel AJ, Secen N, et al. Tiragolumab plus atezolizumab versus placebo plus atezolizumab as a first-line treatment for PD-L1-selected non-small-cell lung cancer (CITYSCAPE): primary and follow-up analyses of a randomised, double-blind, phase 2 study. Lancet Oncol. 2022;23(6):781–92.

45. Rudin CM, Liu SV, Soo RA, Lu S, Hong MH, Lee JS, et al. SKYSCRAPER-02: Tiragolumab in Combination With Atezolizumab Plus Chemotherapy in Untreated Extensive-Stage Small-Cell Lung Cancer. J Clin Oncol. 2024;42(3):324–35.

46. Guan X, Hu R, Choi Y, Srivats S, Nabet BY, Silva J, et al. Anti-TIGIT antibody improves PD-L1 blockade through myeloid and T(reg) cells. Nature. 2024;627(8004):646-55.

47. Rousseau A, Parisi C, Barlesi F. Anti-TIGIT therapies for solid tumors: a systematic review. ESMO Open. 2023;8(2):101184.

48. Li YJ, Du L, Wang J, Vega R, Lee TD, Miao Y, et al. Allosteric Inhibition of Ubiquitin-like Modifications by a Class of Inhibitor of SUMO-Activating Enzyme. Cell Chem Biol. 2019;26(2):278–88 e6.

49. Lv Z, Yuan L, Atkison JH, Williams KM, Vega R, Sessions EH, et al. Molecular mechanism of a covalent allosteric inhibitor of SUMO E1 activating enzyme. Nat Commun. 2018;9(1):5145.

50. Canon JR, A. C, Bradley S, Wang L, Kui M, Wein A, et al. SB-4826, a first-in-class oral, covalent inhibitor of SUMO E1 that induces IFN signaling and inhibits tumor growth as monotherapy and in combination with immune checkpoint blockade. Cancer Res. 2023;83:LB318.

51. Boj SF, Hwang CI, Baker LA, Chio, II, Engle DD, Corbo V, et al. Organoid models of human and mouse ductal pancreatic cancer. Cell. 2015;160(1-2):324–38.

52. Roy I, McAllister DM, Gorse E, Dixon K, Piper CT, Zimmerman NP, et al. Pancreatic Cancer Cell Migration and Metastasis Is Regulated by Chemokine-Biased Agonism and Bioenergetic Signaling. Cancer Res. 2015;75(17):3529–42.

