## Supplementary Figure for "Immunomodulation of Pancreatic Cancer via Inhibition of SUMOylation and CD155/TIGIT Pathway"

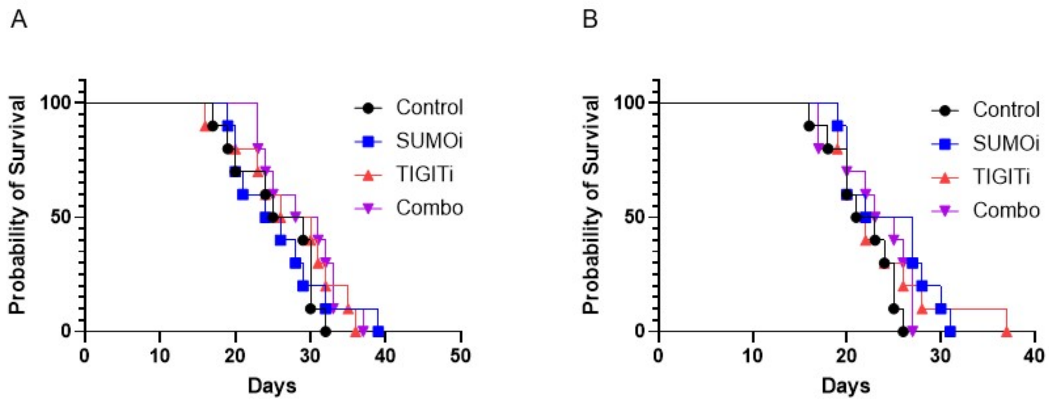


**Supplementary Figure 1.** Kaplan-Meier survival curves of (A) KPC 1199 and (B) KPC 1245 orthotopic pancreatic tumor-bearing C57BL/6 mice treated with SUMOylation and TIGIT inhibitor, alone or in combination.

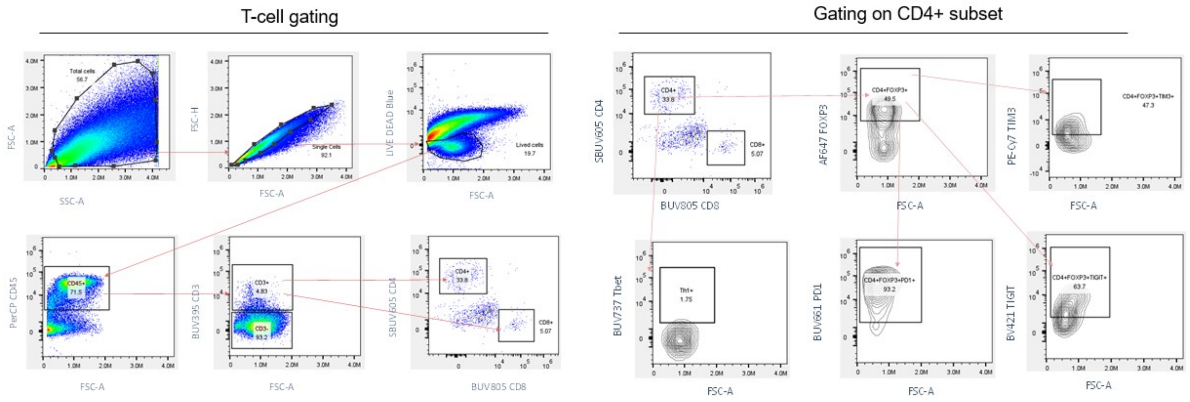


**Supplementary Figure 2**. Flow cytometry gating strategy.
